# Looking for a needle in a haystack: de novo phenotypic target identification reveals Hippo pathway-mediated miR-202 regulation of egg production

**DOI:** 10.1101/2023.10.31.560894

**Authors:** Sarah Janati-Idrissi, Mariana Roza de Abreu, Cervin Guyomar, Fernanda de Mello, Thaovi Nguyen, Nazim Mechkouri, Stéphanie Gay, Jérôme Montfort, Anne Alicia Gonzalez, Marzieh Abbasi, Jérôme Bugeon, Violette Thermes, Hervé Seitz, Julien Bobe

## Abstract

Understanding microRNA (miRNA) functions has been hampered by major difficulties in identifying their biological target(s). Currently, the main limitation is the lack of a suitable strategy to identify biologically relevant targets among a high number of putative targets. Here we provide a proof of concept of successful *de novo* (*i.e*., without prior knowledge of its identity) miRNA phenotypic target (*i.e*., target whose de-repression contributes to the phenotypic outcomes) identification from RNA-seq data. Using the medaka *mir-202* knock-out (KO) model in which inactivation leads to a major organism-level reproductive phenotype, including reduced egg production, we introduced novel criteria including limited fold-change in KO and low interindividual variability in gene expression to reduce the list of 2,853 putative targets to a short list of 5. We selected *tead3b,* a member of the evolutionarily-conserved Hippo pathway, known to regulate ovarian functions, due to its remarkably strong and evolutionarily conserved binding affinity for miR-202-5p. Deleting the miR-202-5p binding site in the 3’ UTR of *tead3b*, but not of other Hippo pathway members *sav1* and *vgll4b*, triggered a reduced egg production phenotype. This is one of the few successful examples of *de novo* functional assignment of a miRNA phenotypic target *in vivo* in vertebrates.

## Introduction

Since the discovery of miRNAs three decades ago, identification of their biological targets has remained a major challenge that has been hampered by the very high number of *in silico*-predicted targets (*i.e*., several hundreds or thousands for a single miRNA) and the high variability in target prediction method outputs (1, 2). In addition, the post-transcriptional action of miRNAs is highly dependent on many factors, including binding affinity of the target site (3, 4), relative abundance of miRNA molecules compared to possible targets expressed in the same cellular context (5), and sequence of the 3’ UTR of the target around the target site, to name just a few. Despite the advent of transcriptomics, quantitative data at the cellular level remain extremely limited to date, at least *in vivo*. For all the above reasons it is therefore extremely difficult to identify biologically relevant targets.

The use of *in vitro* over- or under-expression techniques, even though widely used, suffers from major limitations and has proved to lead to major uncontrolled biases and non-physiological responses (5–7). It is therefore commonly accepted that the most definitive approach for isolating and confirming the importance of a particular target is to disrupt the miRNA target site within the endogenous target gene (8). Such labor-intensive efforts cannot realistically be carried out for hundreds or thousands of putative targets. Currently, the main limitation is therefore the lack of an efficient strategy to identify high-confidence biologically relevant targets among a high number of putative targets. For this reason, a very limited number of functional miRNA/target interactions have been genetically demonstrated *in vivo* (9–13). Two phenotypic targets (*i.e*., targets whose de-repression contributes to the phenotypic outcomes (10)) were identified in nematodes (9, 10), including the let-7/lin-41 interaction, while a single example exists in *Drosophila* (11). In vertebrates, the miR-200/Zeb1 interaction, which controls tumorigenesis in an artificially sensitized mouse model was recently identified (12). In addition, studies in mice have shown that miR-155 could interact with specific genes important for immunity (14–16). Other examples exist that suggest in vivo miRNA/target interaction (17, 18). It should be noted that in most cases the identity of the putative target was already suspected based on prior experimental evidence. In the only case of *de novo* identification (*i.e*., without prior knowledge of its identity), the assignment of the phenotypic target was obtained through a high-throughput reverse genetic screen conducted in *C. elegans*. Collectively, these results illustrate the difficulty of *de novo* identifying biologically relevant miRNA targets, especially in vertebrates in which the possibilities of large-scale genetic screens are limited. They also indicate that a single mRNA target can explain, at least in part, the phenotype under the control of a specific miRNA. It had been speculated that miRNAs control macroscopic phenotypes through the coordinated fine-tuning of multiple genes (19). In contrast with that notion, it therefore appears that, at least in the few cases where miRNA-controlled phenotypes have been precisely dissected, a single key target seems to be chiefly responsible for the biological phenotype. It is therefore not unrealistic to attempt to identify key miRNA targets responsible for organism-level phenotypes.

Despite these limitations, major advances have been achieved in our understanding of miRNA features and mechanisms of action. This includes the report that the amplitude of miRNA-triggered silencing is limited in terms of changes in both mRNA (20, 21) and protein levels (22) (*i.e*., less than 2-fold in most cases). Significant progress has also been made in predicting the efficiency of the miRNA binding site on mRNA target 3’ UTR (3, 4). We reasoned that this knowledge of miRNA function and features as well as in-depth analysis of the phenotype could be used to identify phenotypic targets. As a proof of concept, we used the *mir-202* knock-out (*mir-202* KO) medaka model (23) in which inactivation leads to a major organism-level reproductive phenotype, including reduced egg production. We introduced novel criteria including limited fold-change in gene expression and interindividual variability in wild-types and KO to reduce the list of 2,853 putative targets to a short list of 5. We selected *tead3b,* a member of the evolutionarily-conserved Hippo pathway, known to regulate ovarian functions, due to its remarkably strong and evolutionarily conserved binding affinity for miR-202-5p. Deleting the miR-202-5p binding site in the 3’ UTR of *tead3b* resulted in a phenocopy of the reduced egg production phenotype. This is one of the few successful examples of *de novo* functional assignment of a miRNA phenotypic target *in vivo* in vertebrates. Our work also provides a proof of concept for *bona fide* miRNA target identification.

## Methods

### Establishment of the *mir-202 KO*, *tead3b* [-40]UTR, *sav1* [+14;-46]UTR, *vgll4b* [+26-57]UTR and *vgll4b* [-179]UTR mutant medaka lines

Investigations were conducted in compliance with EU Directive 2010/63/EU on the protection of animals used for scientific purposes and approved by INRAE-LPGP Animal Care and Use Committee under #M-JB012[#1234]. Genome editing was performed using CRISPR/Cas9 to independently delete the genomic regions corresponding to the mir-202 hairpin in the *mir-202* medaka gene (FishmiRNA (24) gene ID FMORYLATG0143, http://fishmirna.org/#) and the miR-202-5p target recognition site in the 3’ UTR region of *tead3b* (ENSORLG00000006370), *sav1* (ENSORLG00000008998), and *vgll4b* (ENSORLG00000004490) genes. Identification of target sites was performed using the CRISPOR tool (http://crispor.org) (25). Several sites were used for each targeted region to obtain genomic deletions. For *mir-202* KO, targets were prepared as previously described (23). For the deletion of the miR-202-5p binding site in *tead3b, sav1,* and *vgll4b* 3’UTR, guide RNAs were prepared as follows. The DR274 plasmid (Addgene # 42250) was used as a template for PCR using a universal primer (AAAAGCACCGACTCGGTGCCACT) and a specific primer fused with the T7 sequence and a sequence corresponding to the plasmid GAAATTAATACGACTCACTATA(*target_genomic_sequence*)GTTTTAGAGCTAGAAATAGCAAG. After digestion of residual plasmid using DpnI for 2-3 h at 37°C, PCR products were purified using a PCR clean-up kit (Macherey Nagel 740609.50). PCR products were subsequently used for guide RNA synthesis using MEGAshortscript T7 Transcription Kit (Invitrogen AM1354) followed by a DNase treatment. Guide RNAs were subsequently phenol-chloroform purified and resuspended in water. For the Cas9 guide RNA, the pCS2-nCas9n plasmid was linearized using Not1 before RNA synthesis. Cas9 RNA was synthesized using the mMESSAGE mMACHINE SP6 Transcription Kit (Invitrogen AM1340). One-cell medaka embryos were injected with 50 ng/ml of each guide RNA, 150 ng/µl of Cas9 RNA, and 200 ng/µl Cas 9 protein (New England BioLabs, M0386T). Injected individuals were screened (see below genotyping procedure) 2-3 months after injection and fish harboring a suitable deletion of the targeted zone were crossed with wild-type individuals to generate a stable line.

### Genotyping of *mir-202* KO, *tead3b* [-40]UTR, *sav1* [+14;-46]UTR, *vgll4b* [+26-57]UTR and *vgll4b* [-179]UTR mutants

Genotyping was performed as previously described (26). Briefly, 2-3 month-old adult individuals were anesthetized using tricaine at 225 mg/l (PharmaQ, tricaine) diluted in water with 450 mg/ml of sodium bicarbonate (Sigma, S5761) and a small piece of the caudal fin was biopsied for genomic DNA extraction. Samples were lysed in 35 µl of lysis buffer containing 1.25 M NaOH and 10 mM EDTA (pH 12) incubated at 95°C for 45 min and the lysis reaction was neutralized by adding 35 µl of neutralization solution containing 2 M Tris-HCl (pH 5). The identification of wild-type and mutant fish was performed by PCR using primers flanking the site of deletion. PCR was performed using JumpStart Taq DNA Polymerase (Sigma-Aldrich, D9307) with the following PCR conditions: 94°C for 2 min, followed by the amplification cycle (94°C, 30 s; 60°C, 30 s; 72°C, 30 s) repeated 40 times. A 2-min final elongation step at 72°C was then performed. The following primers flanking the deletion sites were used for *mir-202* KO (Forward: CGTTCCCAGGAGACATGG, Reverse: CCTCTGTGATGTGAGCAGGA), *tead3b* [-40]UTR (Forward: AGAAGACGTGTGCTTGGATG, Reverse: GCCCCTGTAACTAGTTTGCG), *sav1* [+14;-46] (Forward: CCGTGGCCTGATGAGAAATG, Reverse: CAAGTCCATGCTGCAGGTAC), and *vgll4b* [-179]UTR/*vgll4b* [+26-57]UTR (Forward: CTGGACCCCACAAGATAGCA, Reverse: CCTCTGACGCTTTCCAGTTC) lines.

### Reproductive phenotyping

For all lines, fish generations were obtained through the crossing of heterozygous fish of the previous generation. WT and mutant fish used for phenotyping were siblings of the same age, raised in common at a maximum density of 25 fish in 10-liter tanks and held at 26°C under a (12h light / 12h dark) photoperiod regime for 90 days. Just before the end of this 90-day period, fish were genotyped. At 90 days post-fertilization, fish were then submitted to a specific 15h45 light/ 7h15 dark photoperiod regime that triggered reproduction. Each female was placed in a 1.4-liter tank with a male and monitored individually over the phenotyping period. The phenotyping period began when all WT females started to lay eggs (usually 1 to 2 weeks after photoperiod change). WT males were used for *mir-202* KO. For practical reasons WT and mutant females were mated with males of different genotypes for the *tead3b* [-40]UTR line. The absence of any significant male effect was subsequently verified. For the *mir-202* KO line, 10 mutant and 5 WT females were monitored 2-3 times a week over a 4-week period to confirm the previously described phenotype(23). For the *tead3b* [-40]UTR line, 14 mutant and 10 WT females were monitored 2-3 times a week over a 6-week period. For *sav1* [+14;-46]UTR, *vgll4b* [-179]UTR, and *vgll4b* [+26-57]UTR mutant lines the number of females used is provided in Figure 7. For all lines, spawned eggs were collected from each individual female in the morning after the light was turned on. For each clutch, eggs were counted and classified as developing or non-developing using a stereomicroscope.

To further illustrate the *tead3b*-dependent regulation of egg production by miR-202, the size of the ovary was measured using 90-day-old females in both *mir-202* KO, *tead3b* [-40]UTR lines. Fish were euthanized using an overdose of tricaine at 300 mg/L (PharmaQ, tricaine) diluted in water with 600 mg/ml of sodium bicarbonate (Sigma, S5761). The body cavity was opened and ovaries were fixed *in situ* to preserve their structure. Fixation was performed overnight in 4% paraformaldehyde (PFA) in 1X PBS (pH 7.4) at 4°C under gentle agitation. After fixation, ovaries were dissected out and then rinsed three times in 1X PBS (30 min, 2h, 2h) at room temperature (RT) and subsequently dehydrated in increasing concentrations of ethanol (25, 50, 75, and 100%) diluted in PBS (30 min at RT for each step). Dehydrated ovaries were stored at −20°C in 100% ethanol. Pictures of ovaries were taken on a light table (eSync) using a digital single-lens reflex (SLR) camera (canon EOS 2000D, with an objective EF 100 mm f/2.8 Macro USM). The picture calibration varied between 236.4 and 241 pixels/mm.). The size of the ovaries was calculated using the FIJI software.

### Sampling for RNAscope and miRNAscope

Adult individuals (approximately 3-month-old) were euthanized using an overdose of tricaine as described above. For histological analyses, ovaries were collected from female medaka and fixed overnight in 4% paraformaldehyde (PFA) in 1X PBS (pH 7.4) overnight at 4°C under gentle agitation. After fixation, ovaries were rinsed once in 1X PBS for 10 min at room temperature (RT) and subsequently dehydrated in increasing concentrations of methanol (25, 50, 75, and 100%) diluted in PBS with 0.1% of Tween-20 (10 min at RT for each step). Dehydrated ovaries were stored at −20°C in 100% methanol.

### RNAscope and miRNAscope

RNAscope and miRNAscope procedures were performed using RNAscope 2.5 HD Reagent Kit-RED (ACDbio, 322350) and miRNAscope HD Reagent Kit-RED (ACDbio, 324500), respectively. Most steps are similar for the two protocols except for the steps mentioned in the text. The kits used are different even though some solutions are used in both protocols including Wash Buffer, Target Retrieval, H_2_O_2_ reagent, and the different AMP reagent. The *in situ* hybridization procedure was carried out following the manufacturer’s instructions (ACDBio, Bio-techne, Abingdon, United Kingdom). The RNA scope probes *tead3b* (ACDBio, 1216391-C1) and miRNA-scope probe miR-202-5p (ACDBio, 1127801-S1) were designed by the ACDBio company.

### Ovarian section preparation

Before the RNA or miRNA-scope procedure, fixed ovaries were embedded in paraffin, and sections of 7 µm thickness were obtained using a microtome (microm HM355S, GMI, Ramsey, MN 55303, USA) and placed on adhesive glass slides (VWR, Superfrost Plus). For deparaffinization and rehydration, slides were warmed for 1 h at 60°C and deparaffined in two successive baths of xylene for 5 min at RT, followed by two successive baths of 100% ethanol for 2 min at RT. After deparaffinization, slides were dried at 60°C for 5 min.

### Sample pre-treatment and unmasking

Samples were pre-treated with H_2_O_2_ for 10 min at RT to block endogenous peroxidase activity. The slides were rinsed twice in sterile water. Unmasking was performed by incubating the slides in the 1X RNAscope Target Retrieval reagent solution, previously boiled at 100°C, for 15 min (ACDBio, 322000). This step prevents the cross-linking that occurs from tissue fixation (Bio-techne). The slides were then rinsed twice in sterile water and transferred for 3 min in 100% ethanol before being dried completely at RT. Ovarian sections were permeabilized using protease for 25 min at 40°C (miRNA-scope: Protease III, RNA-scope: Protease plus).

### Probe hybridization

Hybridization was performed for 2h at 40°C using a pre-warmed probe solution.

– miRNA-scope: probe SR-ola-miR-202-5p-S1 (ACDBio, 1127801-S1)
– miRNA-scope: negative control, probe SR-Scramble-S1 (ACDBio, 727881-S1)
– RNA-scope: probe OI-tead3b-C1 (ACDBio, 1216391-C1)
– RNA-scope: negative control, probe-DapB (ACDBio, 310043)

The slides were then washed twice in 1X Wash Buffer for 2 min at RT. After this step, the slides were immersed in 5X SSC solution overnight (O/N) at RT (Euromedex, EU0300-C). The following day, slides were washed twice in 1X wash buffer for 2 min at RT before the signal amplification steps. Six steps were carried out to hybridize preamplifiers and amplifiers, each one with a different reagent. Each step started with adding reagents on ovarian sections. Slides were incubated at 40°C for the first four steps and at RT in the dark for the rest of the protocol. The steps were as follow: AMP1: 30 min, AMP2: 15 min, AMP3: 30 min, AMP4: 15 min, AMP5: 30 min, AMP6: 15 min. Finally, the signal was revealed by adding FastRED-B diluted in FastRED-A (1:60) on ovarian sections, for 15 min at RT in the dark. Slides were then washed twice for 2 min in sterile water. Nuclei were counter-labelled with DAPI (0.1 µg/ml) for 30 s at RT and cover slides were mounted using ProLong Gold antifade reagent (Thermo Fisher Scientific, P36934). Pictures were acquired using a fluorescent microscope (Leica, DM6B) and processed under Fiji software (Image J, release 2.3.0, National Institutes of Health, Bethesda, MD) to adjust the color balance.

### RNA-seq analysis

#### Ovarian follicle sampling

Ovaries from 9 WT and 11 KO *mir-202* medaka were manually dissected using fine forceps. Early (EV) and late vitellogenic (LV) ovarian follicles were isolated based on their diameter (EV: 200-300 µm, LV: 400-500 µm) (27). Pools of 20-30 ovarian follicles originating from the same individual were frozen in liquid nitrogen and stored at −80°C until RNA extraction.

### RNA preparation

RNA extraction was performed as previously described (28). Briefly, frozen follicles were placed in TRI Reagent (Euromedex, Tr118). Samples were subsequently lysed in a Precellys Evolution Homogenizer (Bertin Technologies, Ozyme) in 7 ml Precellys lysing tubes (Bertin Technologies, Tissue homogenizing CK28). Total RNA was subsequently extracted according to the manufacturer’s instructions. After extraction, RNA concentration was measured using a Nanodrop ND-1000 (Nixor Biotech,), and sample integrity was checked using a 2100 Bioanalyser RNA 6000 nano kit (Agilent technologies). All samples presented a RIN value above 8.5. RNA samples were stored at −80°C until library synthesis.

### Library construction and sequencing

All libraries were synthesized from at least 2 µg of template RNA. Polyadenylated RNA was captured using oligo(dT) magnetic beads and libraries were synthesized using Stranded mRNA Prep Ligation (Illumina). RNAs were chemically fragmented and synthesis of the first strand of cDNA was made in the presence of actinomycin D. The second strain of cDNA was synthesized using dUTP. Adapters were ligated to the 5’ end (P5) and 3’ end (P7) and cDNAs were then amplified by PCR. PCR products were purified using AMPure XP Beads (Beckman Coulter Genomics). Libraries were quantified and their quality was checked using Fragment Analyzer NGS High Sensitivity kit (Agilent Technologies) and KAPA Library quantification kit (Roche). Clustering and sequencing were performed on a NovaSeq6000 (Illumina,) using NovaSeq Reagent Kits (Illumina) and the SBS (Sequence By Synthesis) technique. Samples were processed using two lanes of 100-bp single read SP flow cell. More than 1,062 million reads were obtained with a number of reads per library ranging from 18 to 34 million.

### RNAseq data quantification and differential analysis

Raw reads were analyzed using the nf-core/RNAseq Nextflow pipeline in version 3.3 (29). Nf-core is an initiative aiming at providing a set of reference pipelines for bioinformatics analyses. Within the pipeline, raw reads were trimmed, aligned on the Ensembl reference genome in version 104 (accession ASM223467v1), and quantified using Salmon (30). Raw gene counts were processed following the guidelines of the DESeq2 R package in version 1.32.0 (31), using a model including the effects of the development stage (early or late vitellogenesis), the genotype (Wild type or KO), and their interaction. A contrast-based comparison was made to output genes significantly differentially expressed between KO and wild type with an FDR threshold of 0.05.

### Quantitative PCR analysis in the *tead3b* [-40]UTR line

To assess the impact of removing the miR-202-5p binding site in the 3’ UTR region of *tead3b* mRNA levels, ovarian follicles at the EV stage were sampled from 10 WT and 14 mutant *tead3b* [-40]UTR females. Pools of 20-30 ovarian follicles originating from the same individual were frozen in liquid nitrogen and stored at −80°C until RNA extraction. RNA extraction was performed as described above. Quantitative PCR was performed as previously described (23) using the following primer pairs (*tead3b*: GCCATCTACCCTCCATGTGG/ TTTCTTGTGCGGGTTTTGCC, *18S*: CGTTCTTAGTTGGTGGAGCG/AACGCCACTTGTCCCTCTAA). QPCR *tead3b* signal was normalized using *18S*.

### miRNA target prediction

Possible miR-202-5p targets were predicted using Targetscan 7.0 (32, 33). TargetScan was run using the medaka miR-202-5p sequence and the sequences for every feature of the medaka Ensembl annotation with the “three_prime_UTR” value in the feature field.

### Expression variability analysis

The variability of fold changes putatively induced by miRNAs on differentially expressed genes (*i.e*., between WT and KO groups) was compared to the expression variability observed within the wild-type group. Gene expression variability among WT replicates was compared to WT-to-KO differences for every gene by computing the absolute value of every pairwise difference between WT replicates, as well as the absolute value of every difference between a WT replicate and a KO replicate. These differences were calculated from normalized RNA-seq-measured abundances. The set of 36 WT-to-WT differences was then compared to the set of 99 WT-to-KO differences by a one-tailed Wilcoxon test to identify genes with WT-to-KO differences significantly exceeding WT-to-WT differences. Wilcoxon test *P*-values were then corrected using the Benjamini-Hochberg method.

### Comparative analysis of miR-202/target site duplex stability

For each TargetScan-predicted miR-202 target, the predicted binding site was isolated from the whole 3′ UTR (Ensembl Genes 109, medaka genome assembly ASM223467v1) by extracting the sequence spanning 20 nt upstream to 3 nt downstream of the seed match. Its pairing stability with miR-202 was predicted using RNAduplex with default settings.

### Conservation analysis of miR-202/*tead3b* pairing geometry

Orthologs for *O. latipes tead3b* ENSORLG00000006370 were identified in Teleostei using NCBI’s "Homologene" database (https://www.ncbi.nlm.nih.gov/homologene) and the list was manually supplemented with a Holostei sequence (*Lepisosteus oculatus tead3* ENSLOCG00000009736, identified as an ortholog of *O. latipes tead3b* by sequence homology and conserved synteny with the smpd2a/b gene). For each of these 96 species, the longest 3′ UTR isoform among RefSeq mRNA sequences was extracted, resulting in a list of 95 UTR sequences (for species *Chanos chanos*, the only annotated *tead3b* mRNA sequence contains an empty 3′ UTR). UTRs were aligned using ClustalW, and sequences orthologous to the *O. latipes* miR-202 binding site were extracted from that alignment. Mature miR-202 sequence in each species was inferred from aligned mir-202 pre-miRNA hairpins found by a genome-wide homology search or from the FishmiRNA database (24). Sequences orthologous to the *O. latipes* miR-202 binding site in *tead3b*, and species-specific mature miR-202 sequences, were then conceptually annealed using RNAduplex in order to identify their most stable pairing geometry.

### Script and data availability

Scripts, raw and intermediate data used for the identification of candidate targets exhibiting higher WT-to-KO variability than intra-WT variability, for the comparative analysis of miR-202/target site duplex stability, and for the conservation analysis of miR-202/*tead3*b pairing geometry, are available at https://github.com/HKeyHKey/Janati-Idrissi_et_al_2023 and at https://www.igh.cnrs.fr/en/research/departments/genetics-development/systemic-impact-of-small-regulatory-rnas#programmes-informatiques.

## Results

### *miR-202* knock-out phenotype

In a previous study, a small indel in the miR-202-3p sequence resulted in the total absence of both −5p and −3p mature forms and led to a major organism-level organism phenotype, including a dramatic reduction in egg production(23). To further confirm these results and rule out any possible miR-202 expression in KO females, we have used two different guides to obtain a full deletion of the *mir-202* hairpin including the genomic regions corresponding to miR-202-5p and 3p (Figure 1A). To demonstrate the full repression of miR-202-5p, which is the active miR-202 form in fish, we used miRNAscope, a sensitive RNA in situ hybridization procedure specifically dedicated to the visualization of miRNAs, on ovarian sections of wild-type (WT) and *mir-202* KO (KO). On KO sections, we observed a full abolishment of the miR-202-5p signal that was observed in the granulosa cells of WT fish (Figure 1B). Because miR-202 is a so-called “gonad-specific” miRNA almost exclusively expressed in the ovary in females(23), the reproductive phenotype resulting from its inactivation was investigated. In agreement with the phenotypes that we had observed in the miR-202-3p indel mutant (23), we observed a significant decrease in the overall egg production (t-test, *P* = 0.009) (Figure 1C) and in egg developmental success rate (Wilcoxon test, *P* = 1.83e-10) (Figure 1D) in the KO group in comparison to the WT group. When analyzing individual egg production performance between WT (*n* = 5) and KO (*n* = 10) we observed, in addition to the decrease in egg production, a higher inter-individual variability in the KO group with a coefficient of variation (CV) of 46%, in comparison to the WT group (CV = 29%) (Figure 1C). We also observed a significantly reduced ovary size in mutant females in comparison to control WT (Figure 1E).

**Figure 1.**
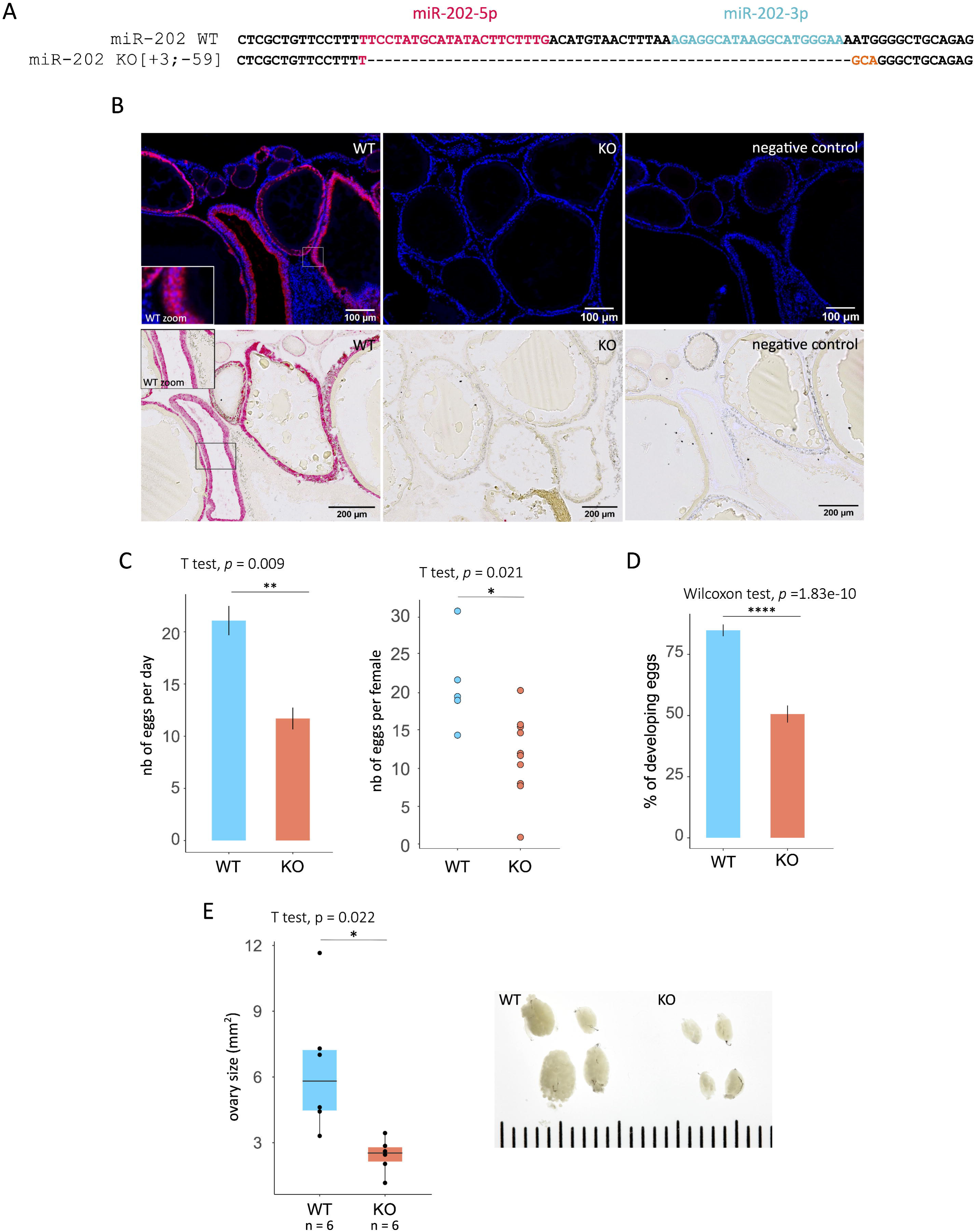
*mir-202* knock out (KO) reproductive phenotype. **(A)** Schematic representation of the [-59;+3] bp indel generated by CRISPR-Cas9 in the *mir-202* gene. Targeted PAM sequences are underlined. **(B)** Fluorescent and chromogenic *in situ* miRNAscope localization of miR-202-5p in wild-type (WT) and *mir-202* knock out (KO) ovaries. **(C)** Mean number of eggs laid per day (histogram) (*n* = 10 days) and mean number of eggs laid per female over the phenotyping period (scatter plot) (WT, *n* = 5; KO, *n* = 10) in WT and KO groups. **(D)** Embryonic development rate of eggs per clutch in WT and KO females (*n* = 10 days). **(E)** Size of the ovary in WT (*n* = 6) and *mir-202* KO mutant (*n* = 6) 90-day-old females (left panel). Representative picture of WT and *mir-202* KO mutant ovaries (right panel). Statistical tests and their *P*-values are indicated above each graph and the degree of significance is represented by asterisks. *P*-values \**P* < 0.05, \*\**P* < 0.001, \*\*\**P* < 0.0001 and \*\*\*\**P* < 0.00001.

### RNA-seq analysis of ovarian follicles in WT and *mir-202* KO fish

To determine the impact of mir-202 KO on gene expression, we dissected individual ovarian follicles from whole ovaries originating from WT and KO groups at early vitellogenic (EV) and late vitellogenic (LV) stages. To standardize ovarian follicular development, ovaries were taken at the same time during the reproductive cycle following ovulation, and ovarian follicles were individualized based on their size. We generated RNA-seq data originating from ovarian follicles of both groups using a high number of replicates (n= 9 in WT, n = 11 in KO). Differentially expressed genes (DEGs) in EV (Table S1) and LV (Table S2) groups (corrected P value <0.05) are shown in Figure 2, including DEGs common to both stages.

**Figure 2.**
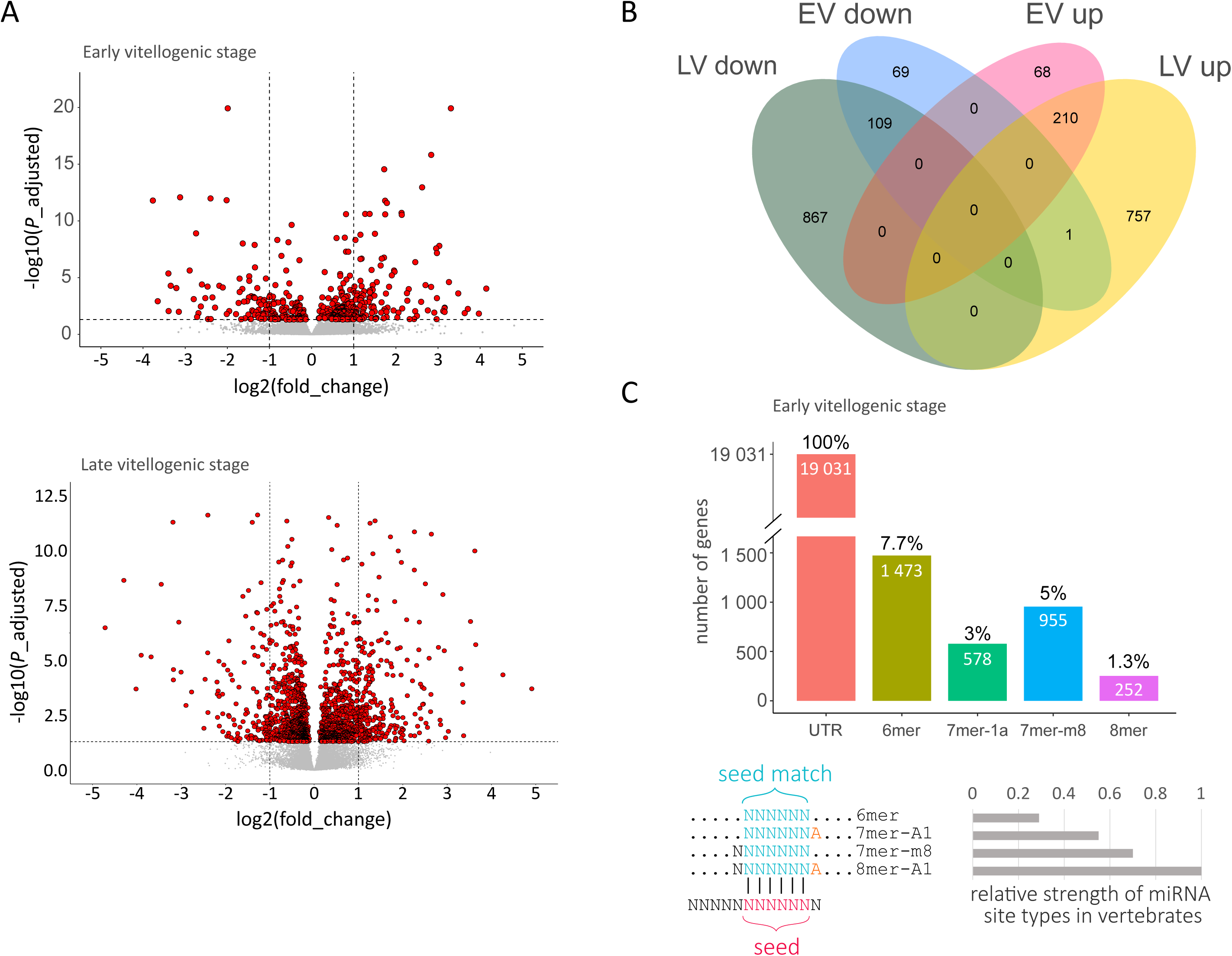
RNA-seq analysis of isolated ovarian follicles from wild-type (WT) and *mir-202* KO (KO) fish. **(A)** Volcano plot representing differentially expressed genes (DEGs) at early vitellogenic (EV) and late vitellogenesis (LV) stages. **(B)** Venn diagram displaying up- and down-regulated DEGs at EV and LV stages, including common ones. **(C)** Proportion of 6mer, 7mer-1a, 7mer-m8, and 8mer predicted targets among expressed genes at EV stages according to TargetScan. The number of genes displaying at least a putative target site in their 3’ UTR region is indicated for each category, including genes that also belong to other categories. The percentage of genes falling into the different categories was calculated relative to the total number of expressed genes (n= 19,031).

Using the TargetScan software, we were able to identify 2,853 putative targets of miR-202-5p among the 19,031 expressed genes that we could measure above background levels at the EV stage and had 3’ UTR sequence information (Figure 2C). The relative efficacy of miRNA repression depends on the binding of the miRNA on a sequence matching the seed of the miRNA in the 3’ UTR region of targeted genes (8). This seed-matched sequence is composed of 6 nucleotides (nt 2 to 7), and additional nucleotides surrounding the seed-matched region. If there is an A facing miRNA position 1 (7mer-A1) efficacy is enhanced. If there is a match on the 8^th^ position (7mer-m8) then the targeting efficacy is even higher. The highest efficacy is observed when both features are present (8-mer) (Figure 2C) (8). The percentage of expressed genes in which at least a site of the above-listed categories can be found is displayed in Figure 2C.

### Novel strategy to identify candidate phenotypic target(s)

Because differences in oocyte content are already observed at the early-vitellogenic (EV) stage in *mir-202* KO fish(23), we reasoned that the expression of miR-202-5p targets regulating egg production was likely to be modified in EV follicles. Using this criterion, we obtained a list of 80 genes (Figure 3A) (Table S1) that were both differentially expressed in EV follicles and *in silico* predicted to bind miR-202-5p in their 3’ UTR region.

**Figure 3.**
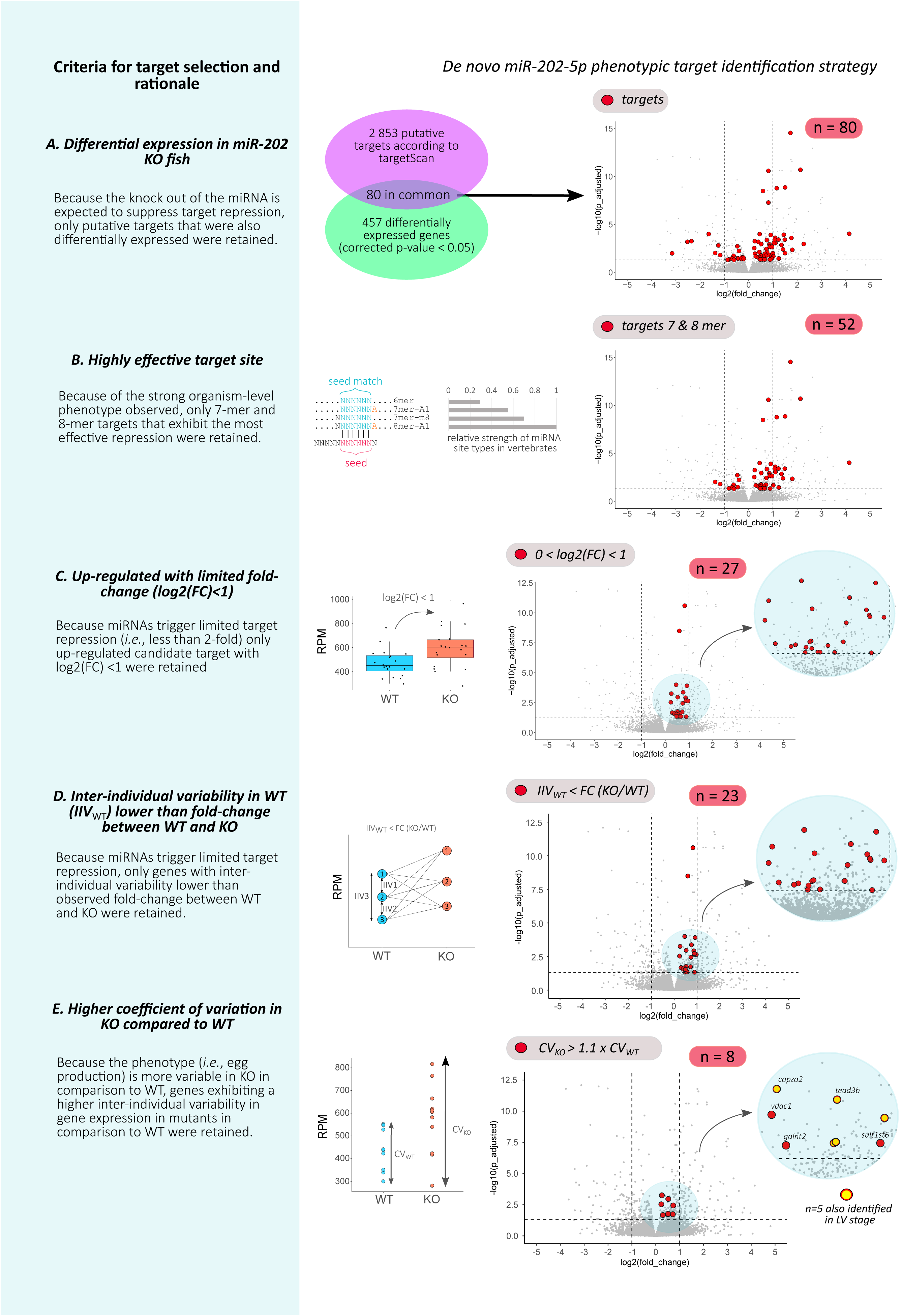
*De novo* miR-202-5p phenotypic target identification strategy. For the different steps, the rationale is presented in the left panel, schematically illustrated in the center panel, and selected genes are shown as red dots on volcano plots in the right panel. **(A)** Selection of genes that are both differentially expressed and putative miR-202-5p targets according to TargetScan. **(B)** Selection of 7mer and 8mer targets among genes identified in A. **(C)** Selection of genes exhibiting a limited overexpression in the KO group (log2FC < 1) among genes identified in B. **(D)** Selection of genes exhibited a limited inter-individual variability in the WT group among genes identified in C. **(E)** Selection of genes exhibited a greater coefficient of variation (CV) in the WT group than in the KO group among genes identified in D.

Because the most effective canonical sites are 7–8 nt sites that include a perfect match to the miRNA seed (positions 2–7 of the seed) (8), only 7mer-A1, 7mer-m8, and 8mer sites were considered for further analysis. In theory, multiple low-efficacy sites (like 6mer sites) may be collectively as efficient as a single high-efficacy site (like a 7 or 8mer site). However, 6mer sites are not only less efficient in guiding gene repression, they are also less conserved in evolution (34), indicating that a single 7mer or 8mer site is better maintained in evolution than a collective set of 6mer sites. We therefore only considered 7mer and 8mer sites. Implementation of this criterion in the phenotypic target identification pipeline led to a reduced list of 52 putative target genes (Figure 3B) (Table S1).

MicroRNAs are known to trigger limited differences in mRNA levels (19, 20) and sometimes lead to differences in protein levels with very limited changes in mRNA levels (20, 21). The first identified miRNA phenotypic target in *Caenorhabditis elegans* was found to be translationally repressed whereas target mRNA levels were only mildly down-regulated, with a log2FC of approximately 1 (9). An earlier study had shown that overexpressing a miRNA in human cell lines causes mostly mild changes in mRNA levels (less than two-fold) (35). In a cellular model, a genome-wide study reported that the loss of miR-223 resulted in mild differences in gene expression with log2FC almost systematically lower than 1 (20). Similarly, genome-wide analysis of mRNA down-regulation following miRNA overexpression resulted in mild responses with log2FC almost systematically below 1 (21). Based on this experimental evidence, we reasoned that changes in gene expression triggered by miR-202-5p in the ovarian follicle were likely to be modest. In addition, because miRNA are post-transcriptional regulators leading to mRNA decay or translational repression(8), mostly through de-adenylation, we hypothesized that miR-202-5p targets were likely to be up-regulated in *mir-202* KO follicles (*i.e*., in the absence of miR-202). For these reasons, DEG were filtered for up-regulated genes in *mir-202* KO individuals with limited fold change (*i.e*., 0<log2FC<1). Implementation of this criterion in the target identification pipeline led to a reduced list of 27 putative target genes (Figure 3C) (Table S1).

We then reasoned that, because miRNAs trigger limited changes in mRNA levels, the inter-individual variability among wild-type samples was likely to be modest. Indeed, a specific miRNA is unlikely to trigger modest changes in mRNA levels if the observed differences in expression levels in WT are high (*i.e*., higher than the maximum expected difference in gene expression in KO compared to WT). We took advantage of the large number of biological replicates (WT, *n* = 9; *mir-202* KO, *n* = 11) to calculate the difference in expression among WT samples. This inter-individual variability in expression among WT was then compared to all possible differences in gene expression between WT and *mir-202* KO individuals. Only genes exhibiting a significantly lower inter-individual variability in WT than the observed FC between WT and *mir-202* KO groups with a corrected *P*-value threshold of 0.05 were considered as credible targets. Implementation of this criterion in the target identification pipeline led to a reduced list of 23 putative target genes (Figure 3D) (Table S1).

To further narrow down the list of putative targets, we went back to the phenotype and noticed that the number of eggs in *mir-202* KO fish was not only lower than in WT but also more variable (Figure 1C) with a coefficient of variation (CV) of 46% and 29% in *mir-202* KO and WT, respectively. We reasoned that the phenotype was under the control of miR-202 direct target(s) and hypothesized that the CV in target gene expression was also likely to be higher in KO fish compared to WT. Implementation of this criterion in the target identification pipeline led to a reduced list of 8 putative target genes (Figure 3E) (Table S1). Among these genes were *tead3b*, *capza2*, *vdac1*, *galnt2*, *sult1st6*, and 3 unannotated genes. In this short list, *tead3b* immediately caught our attention. Tead directly interacts with Yap/Taz, which are the main effectors of the Hippo Pathway, an evolutionarily conserved pathway known to integrate mechanical signals to regulate organ growth and size (36–39). The Hippo pathway components play important roles in follicle growth and activation, and in the maintenance of fertility (40).

As presented here, the number of DEGs at the LV stage (*n* = 1,943) is much higher than at the EV stage (*n* = 456). While a significant proportion of DEGs at the LV stage are possibly the indirect consequence of earlier dysregulations, we reasoned that miR-202-5p targets would also be repressed at the LV stage. To test this hypothesis, we carried out the exact same procedure at the LV stage (Table S2) and identified 5 genes common to both stages. Among these genes were *tead3b*, *capza2*, and 3 unannotated genes (Figure 3E). This further strengthened our choice of selecting *tead3b* for further functional analysis.

#### The miR-202-5p/*tead3b* physical interaction is remarkable and evolutionarily conserved

To assess the ability of miR-202-5p to bind *tead3b* 3’UTR and further evaluate its credibility as a miR-202-5p phenotypic target, we thoroughly characterized the ability of miR-202-5p to bind its target site located in tead3b 3’ UTR (Figure 4A). Using RNAduplex (41), we were able to computationally predict the binding affinity of miR-202 for its target site in *tead3b* 3’UTR. We observed a very strong interaction with a predicted binding (ΔG) of −14.90 kcal.mol^-1^ (Figure 4A). This predicted binding affinity for *tead3b* was higher than for 95% of the 3,733 sites (in 2,853 3’ UTRs) found among expressed genes at the EV stage. Similarly, miR-202-5p binding affinity for *tead3b* was higher than for 96% of the 1,081 7mer-m8 sites (955 genes) found among expressed genes at the EV stage (Figure 4B). Similar results were obtained when the analysis for restricted to the 80 differentially expressed genes between WT and KO (Figure 4C). We then compared the ability of the predicted miR-202-5p binding site in *tead3b* to bind miR-202-5p, to those located in the other candidate genes identified at the EV stage. Overall, these 8 genes exhibit 17 binding sites for miR-202-5p. We observed that only one site among 17 exhibited a higher affinity for miR-202-5p. This 7mer-1A site was found in the 3’UTR of the unannotated gene ENSORLG00000026596. Because this predicted 3’UTR region was surprisingly long when compared to orthologous genes in closely related species and not consistently annotated by NCBI and Ensembl, this site was not retained for further analysis.

**Figure 4.**
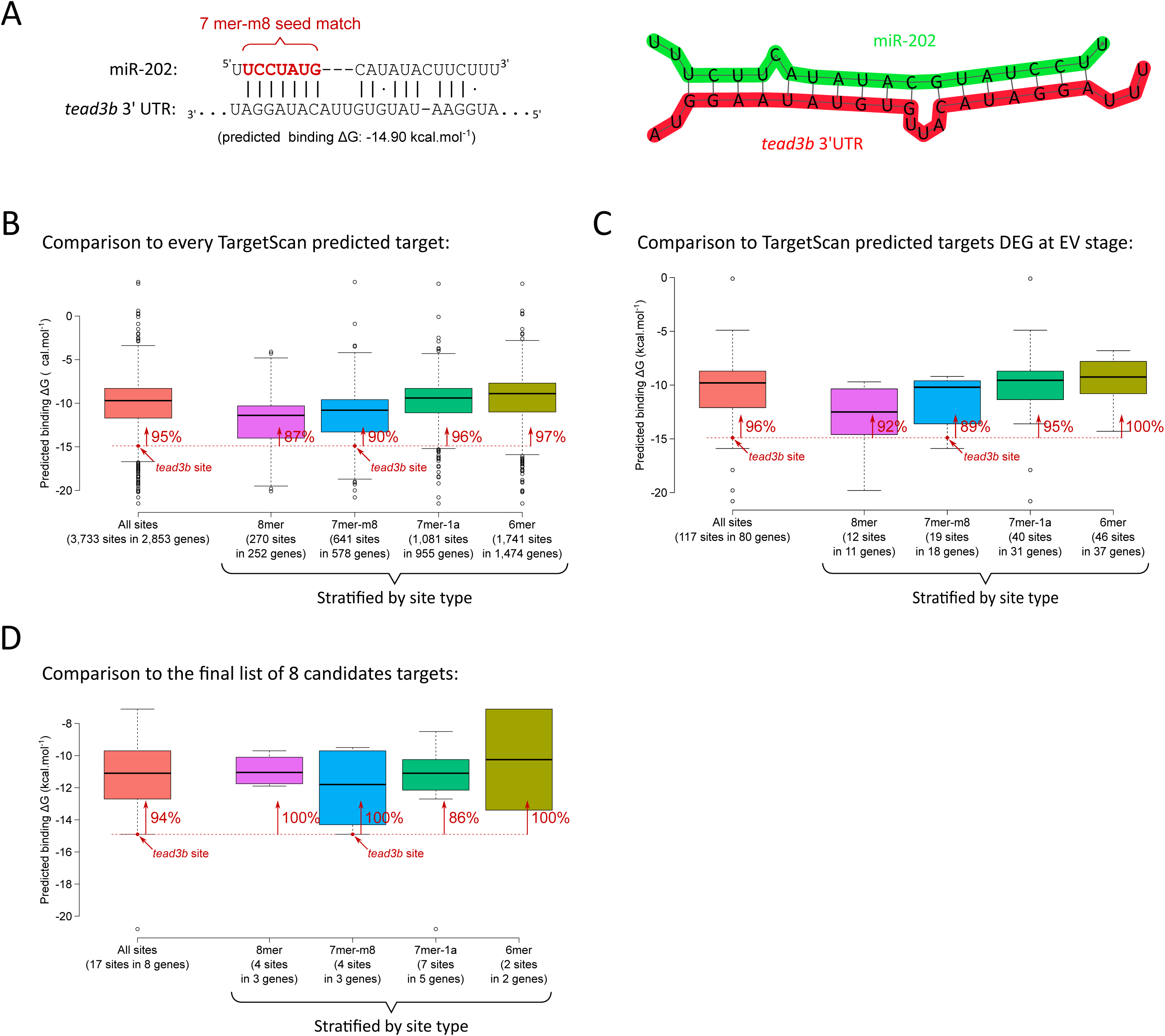
Comparison of predicted miR-202/mRNA duplex stability across predicted miR-202 targets. **(A)** predicted pairing between miR-202 and the *tead3b* 3′ UTR according to RNAduplex (with predicted folding free energy ΔG). The seed match is highlighted in red. **(B-D)** comparison to the predicted ΔG for TargetScan-predicted miR-202 targets in *O. latipe*s, for various sets of candidate targets. In each boxplot, the leftmost box represents every predicted target in that set (regardless of seed match type), while the 4 rightmost boxes are stratified by seed match type. The red dot represents the miR-202 binding site in *tead3b*, and red arrows indicate the percentage of targets whose interaction with miR-202-5p is predicted to be less stable than the *tead3b*/miR-202 interaction.

To further support the relevance of selecting *tead3b* as a miR-202-5p phenotypic target, we took an evolutionary perspective and examined the binding affinity of miR-202 for *tead3b* 3’ UTR in 94 teleost species and 1 holostean species, the spotted gar (*Lepisosteus oculatus*). We observed a strong conservation of the high miR-202-5p/*tead3b* sequence complementarity in the Ovalentaria clade (21 species, including medaka) (Figure 5). Similarly, we observed a high sequence complementarity between miR-202-5p and its target site in different percomorph clades including Ovalentaria, Anabantaria, Carangaria, Gobiaria, Pelagiaria, Syngnatharia, and Eupercaria. In contrast, we observed a much weaker sequence complementarity between miR-202-5p and *tead3b* in other clades including Protacanthopterygii and Otocephala (Clupeomorpha, Characiphysae, Siluriphysae, and Cypriniphysae). However, a strong miR-202/*tead3b* sequence complementarity was observed in spotted gar, a species belonging to the Holostean clade. Teleostean and Holostean lineages have diverged approximately 300 Mya(42). We hypothesize that the ability of miR-202-5p to bind *tead3b* 3’UTR was present in the last common ancestor of Holosteans and Teleosts and was subsequently independently lost in the Otocephala and Protacanthopterygii lineages.

**Figure 5.**
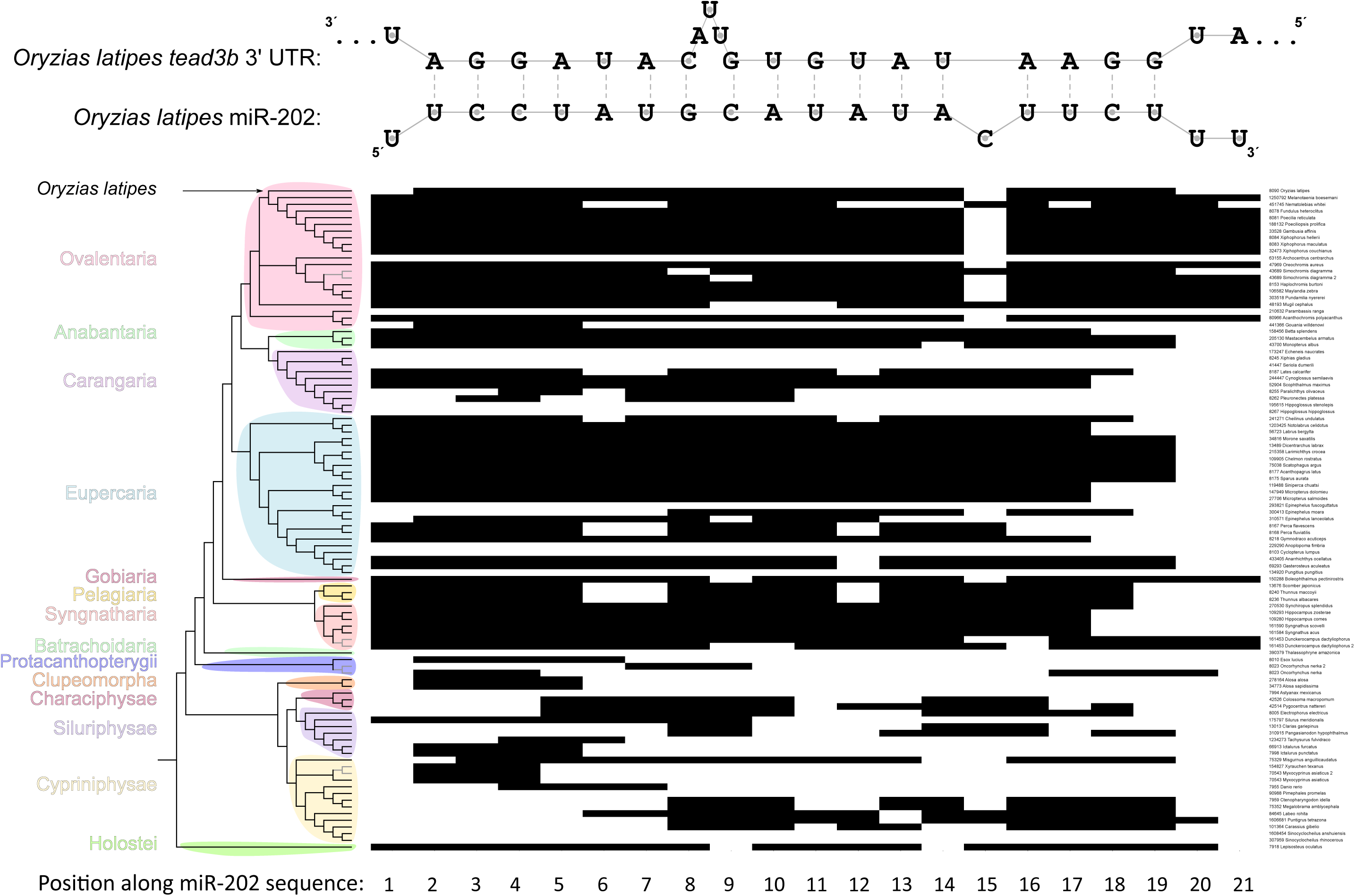
Conservation of miR-202/*tead3*b pairing geometry among Neopterygii (miRNA-centric view; see Figure S1 for target-centric view of the duplex). Each column in the heatmap represents a miR-202 nucleotide (from nt 1 to 21),and each row represents one species (phylogenetic cladogram shown on the left margin). Four species possess genes expressing two distinct mature miR-202 sequences: for these species, two alternative pairing geometries are therefore possible between miR-202 and the site in *tead3b* 3′ UTR; they are represented by two rows on the heatmap, and linked by a grey fork in the cladogram. In the heatmap, a black rectangle denotes a paired nucleotide (either a standard Watson-Crick pair, or a wobble G-U pair), and a white rectangle denotes an unpaired nucleotide. *O. latipes* miR-202 and target site sequences are shown at the top for reference.

### Validation of *tead3b* as a miR-202-5p phenotypic target regulating egg production

#### Both *mir-202* knock out and deletion of the miR-202-5p binding site in *tead3b* 3’UTR lead to significantly reduced egg production

We decided to use *tead3b* for functional analysis due to its key role in the evolutionarily-conserved Hippo signaling pathway known to regulate ovarian follicle growth in vertebrates (40) and its remarkably strong and evolutionarily conserved binding affinity for miR-202-5p. Using CRISPR/Cas9, we deleted a short genomic region of 40 bp containing the miR-202-5p binding site of the 3’ UTR region of *tead3b* (Figure 6A). Deleting the miR-202-5p target site in *tead3b* 3’UTR resulted in a marked reduction in egg production in comparison to control fish (T-test, p = 0.00017, F3 generation) (Figure 6B) and similarly to the phenotype observed in *mir-202* KO females. The average number of eggs per clutch measured over the phenotyping period ranged from 16 to 31 in the WT group. In contrast, the number of eggs per clutch never exceeded 22 in the *tead3b* [-40]UTR mutant group, with a majority of females producing less than 16 eggs per clutch, on average (Figure 6B). The overall difference in egg production is highly significant despite the natural variation observed in WT. In addition, we observed a dramatic decrease in the size of the ovary in the *tead3b* [-40]UTR mutant group in comparison to WT (Figure 6D) in 90-day-old females that we also observed in *mir-202* KO fish (Figure 1E). Together, our observations clearly show that deleting the miR-202-5p binding site in the 3’UTR region of tead3b has major consequences on ovarian development that are similar to those observed in *mir-202* KO fish and ultimately result in a reduced egg production phenotype.

**Figure 6.**
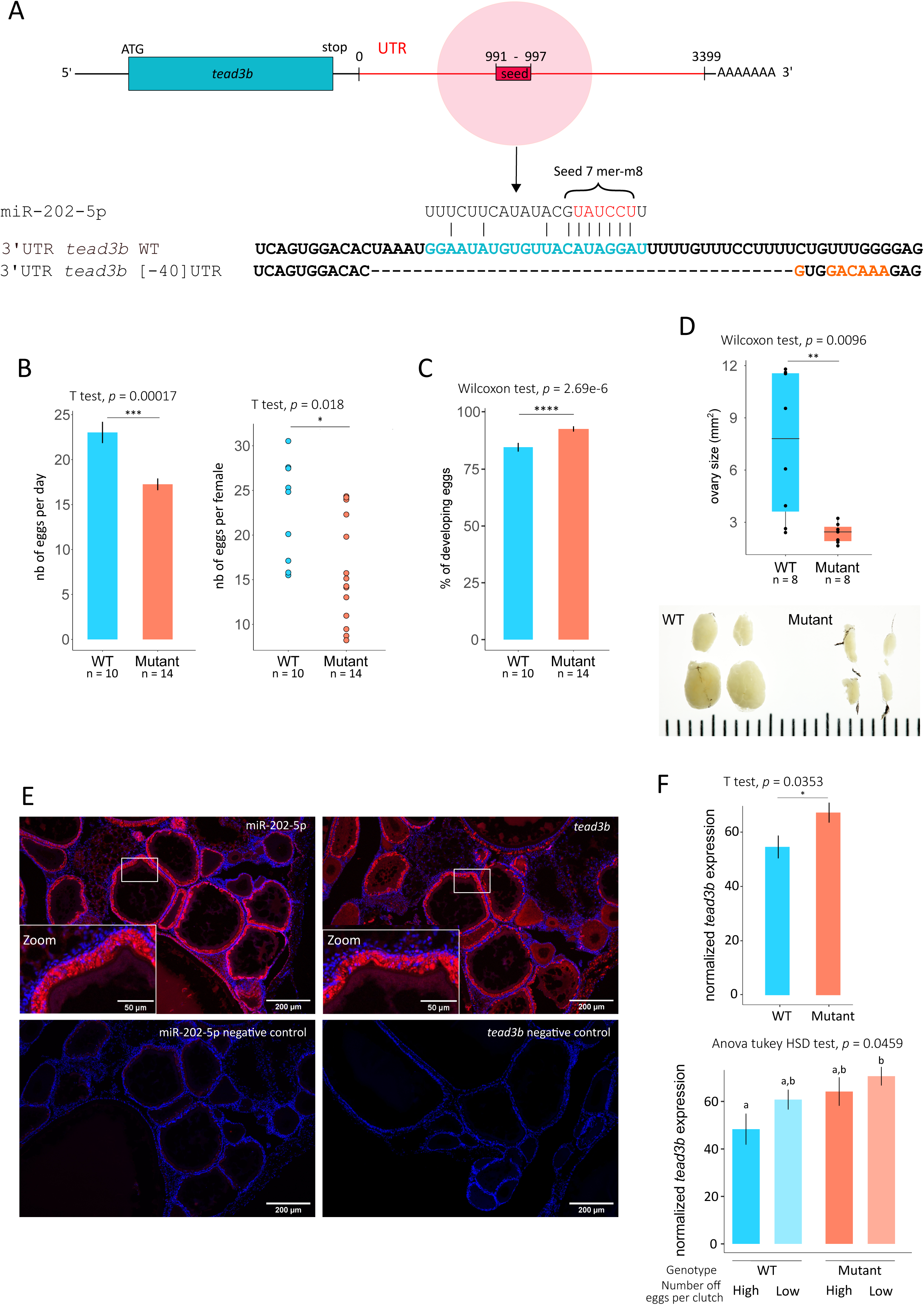
Validation of *tead3b* as the miR-202-5p phenotypic target regulating egg production. **(A)** Schematic representation of the [-40]bp deletion of the miR-202 seed match sequence, generated by CRISPR-Cas9 in the 3’ UTR of *tead3b*. **(B)** Mean number of eggs laid per day (histogram) (*n*=16 days) and mean number of eggs laid per female (scatter plot) (WT, *n* = 10; *tead3b* [-40]UTR mutant, *n* = 14) in WT and mutant fish. F3 fish were used. **(C)** Developmental success of eggs in WT and *tead3b* [-40]UTR mutant females (*n* = 16 days). F3 fish were used. **(D)** Size of the ovary in WT (*n* = 8) and *tead3b* [-40]UTR mutant (*n* = 8) 90-day-old females (upper panel). Representative picture of WT and *tead3b* [-40]UTR mutant ovaries (lower panel). F4 fish were used. **(E)** Fluorescent *in situ* RNAscope localization of *tead3b* in WT ovaries. **(F**) QPCR analysis of *tead3b* mRNA levels in ovarian follicles of control (WT) and *tead3b* [-40]UTR mutant ovarian follicles at EV stage. Statistical tests and their *P-*values were indicated above each graph and the degree of significance was represented by asterisks. *P*-values \**P* < 0.05, \*\**P* < 0.001, \*\*\**P* < 0.0001 and \*\*\*\**P* < 0.00001.

**Figure 7.**
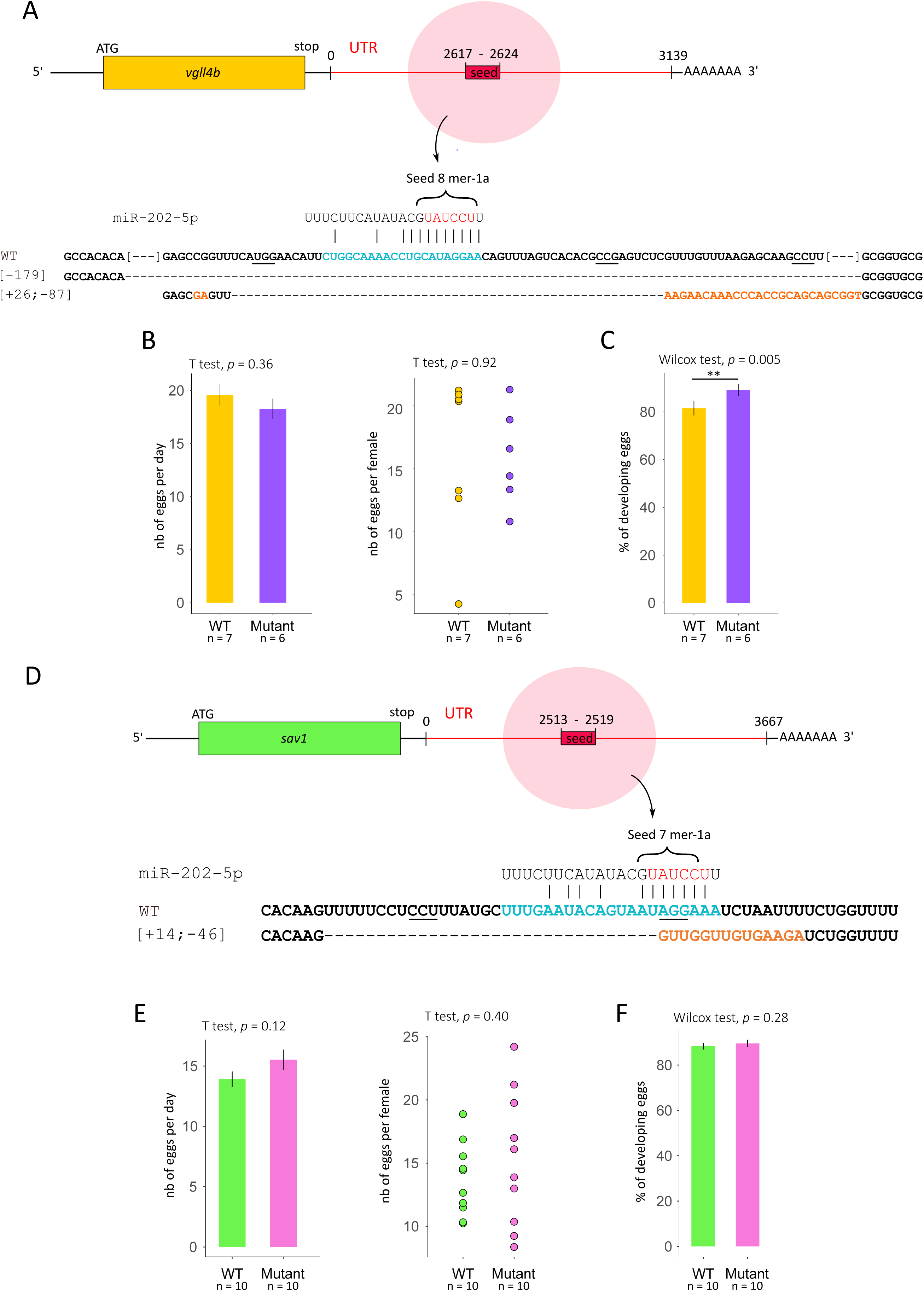
Assessment of *vgll4b* and *sav1* as possible miR-202-5p phenotypic targets regulating egg production. **(A)** Schematic representation of −179bp and +26-57bp indels around the miR-202 seed match sequence, generated by CRISPR-Cas9 in the 3’ UTR of *vgll4b*. **(B)** Mean number of eggs laid per day (histogram) (*n*=13 days) and mean number of eggs laid per female (scatter plot) (WT, *n* = 7; *vgll4b* [+26-156]UTR mutant, *n* = 4; *vgll4b* [-179]UTR mutant, *n* = 2) in WT and mutant fish. F2 fish were used. **(C)** Developmental success of eggs in WT and *vgll4b* UTR mutant females (*n* = 16 days). F2 fish were used. **(D)** Schematic representation of +14-46bp indels around the miR-202 seed match sequence, generated by CRISPR-Cas9 in the 3’ UTR of *sav1*. **(E)** Mean number of eggs laid per day (histogram) (*n*=9 days) and mean number of eggs laid per female (scatter plot) (WT, *n* = 10; *sav1* [+14-46]UTR mutant, *n* = 10) in WT and mutant fish. F2 fish were used. **(F)** Developmental success of eggs in WT and *sav1* UTR mutant females (*n* = 16 days). F2 fish were used. Statistical tests and their *P-*values were indicated above each graph and the degree of significance was represented by asterisks. *P*-values \**P* < 0.05, \*\**P* < 0.001, \*\*\**P* < 0.0001 and \*\*\*\**P* < 0.00001.

In contrast to *mir-202* KO fish (Figure 1D), *tead3b* [-40]UTR mutant fish did not show a decrease in egg developmental success rate suggesting that the two phenotypes (*i.e*., egg number and egg developmental competence) are independently regulated by different miR-202 targets (Figure 6C). Surprisingly, a limited, yet significant, increase in the percentage of developing eggs was observed in the mutant group in comparison to WT.

To further support our observations, we used a similar genome editing approach to delete the miR-202-5p binding site in the 3’UTR regions of two other members of the Hippo pathway and used them as negative controls. We selected *vgll4b*, a downstream player that directly interacts with *tead*, and *sav1* which is the direct partner of mst1/2 (*i.e.*, the vertebrate ortholog of Hippo) acting at the core of the Hippo pathway (39). For both *sav1* and *vgll4b* (Figure 7), deleting the miR-202-5p binding site in the 3’UTR did not have any significant impact on egg production. Together, our observations suggest that miR-202 is acting on the Hippo pathway through *tead3b* but not *sav1* or *vgll4b* to regulate egg production.

#### miR-202 and tead3b are colocalized in granulosa cells in ovarian follicles

Using *in situ* hybridization, we showed that *tead3b* was detected in the ovary with a strong expression in granulosa cells (Figure 6D) in WT fish, similarly to miR-202-5p (Figure 1B). Using adjacent sections, we observed that miR-202-5p and *tead3b* exhibit a strikingly similar expression pattern and are colocalized in granulosa cells within the ovary, including in EV follicles (Figure 6D).

#### Both *mir-202* knock out and deletion of the miR-202-5p binding site in *tead3b* 3’UTR lead to *tead3b* de-repression in ovarian follicles

To further support the association between miR-202-5p and *tead3b*, we monitored *tead3b* mRNA levels in EV follicles in *tead3b* [-40]UTR mutant fish under the hypothesis that the absence of miR-202-5p binding site would result in an increased abundance of *tead3b* mRNA levels. We observed a limited, yet significant, increase in *tead3b* mRNA levels in EV ovarian follicles when the miR-202-5p binding site was removed from the 3’UTR region (Figure 6F). The observed fold-change is fully consistent with expected limited changes in expression levels classically triggered by miRNAs (1, 20, 21). For each genotype, individuals were separated into two groups based on their egg production phenotype using the median in each group as a cut-off. We observed a possible direct link between egg production and *tead3b* mRNA level, even though the difference was only significant between good spawner WT and bad spawner mutants. However, the statistical power of the analysis was limited due to the number of individuals in each group.

## Discussion

### Phenotypic target identification reveals Hippo pathway-mediated miR-202 regulation of egg production

It is currently admitted that “the most definitive approach for isolating and confirming the importance of a particular target is to disrupt the miRNA-binding site(s) within the endogenous target gene and examine the extent to which the resulting de-repression of that target phenocopies the miRNA knockout” (8). In the present study, we used this gold standard genetic approach to demonstrate that miR-202-5p is regulating egg production through *tead3b*. The region that we deleted or modified in the 3’ UTR region of *tead3b* 3’ UTR is very small (40 bp, including 22 bp corresponding to miR-202-5p binding). It is therefore unlikely that the observed phenotype is not due to the lack of miR-202-5p binding in *tead3b* 3’ UTR. In addition, we showed that both miR-202-5p and *tead3b* are both expressed in the granulosa cells within the ovary. We also independently report a limited overexpression (log2FC<1) of tead3b when miR-202 was absent or when the miR-202-5p binding site was removed from *tead3b* 3’ UTR. These results are in favor of a direct regulation of *tead3b* mRNA levels by miR-202-5p. Together our observations strongly suggest that *tead3b* is a miR-202-5p phenotypic target regulating egg production. However, it is possible that other miR-202-5p targets, that remain to be identified, also contribute to the regulation of egg production. Tead3b belongs to the Hippo pathway, an evolutionarily conserved molecular signaling pathways known for its role in regulating organ size and growth (36, 38–40). In mammals, recent evidence has shown that the Hippo pathway plays an important role in regulating reproductive physiology, including the regulation of follicle growth (40). However, the regulation of the Hippo pathway by miR-202 was never reported in any animal species. In the fish ovary, the role of the Hippo pathway in regulating follicle growth or egg production was never investigated but recent studies have shown its importance in micropyle formation in zebrafish (43, 44). Our results are consistent with recent data obtained in mammals and suggest an important role of Hippo signaling in regulating egg production under miR-202 control. Here we report that deleting miR-202-5p binding site in 3’UTR of *sav1* and *vgll4b* (two other members of the Hippo pathway) did not yield any significant defect in egg production, in contrast to *tead3b*. Together, our observations show that miR-202 is targeting Hippo signaling to regulate egg production through its interaction with *tead3b*, and apparently not through other Hippo pathway members *sav1* and *vgll4b*. In addition, the dramatic reduction in the size of the ovary in mutants lacking miR-202-5p binding site in *tead3b* 3’UTR is highly consistent with the role of the Hippo pathway in regulating organ size and suggests a *tead3b*-dependent miR-202 regulation of ovarian development.

### Conservation of the miR-202-5p/tead3b binding affinity during evolution

We observed a remarkable affinity between miR-202-5p and *tead3b*. This high sequence complementarity is not limited to the seed region and is observed over the entire miRNA sequence. This high affinity is not only observed in closely related species but also conserved among a majority of Percomorphs. In other Teleost clades, the interaction appears to be mostly lost, even though present in some species. More importantly, the high affinity of miR-202-5p for *tead3* is found in the gar, a slow-evolving species (45) belonging to a lineage that diverged from Teleosts 300 Mya (42). These observations are in favor of an ancient interaction between miR-202-5p and *tead3b* that was subsequently independently lost in some specific species or clades. The present analysis suffers from methodological limitations, including genome quality, *tead3b* ortholog identification, and quality of 3’ UTR sequence prediction. It is therefore possible that the miR-202-5p/*tead3b* interaction was missed in our analysis, at least for some species. It is also possible that miR-202-5p could target, or at least exhibit sequence complementarity, with other *tead* genes in teleost fish. Interestingly, miR-202-5p exhibits two predicted target sites (8mer and 7mer-m8) in the 3’ UTR of *Tead1* (ENSMUST00000059768.11) in mice, according to TargetScan (https://www.targetscan.org/mmu_80/) (data not shown), while no target site is described in *tead3* (ENST00000402886.3). A comprehensive analysis of mir-202-5p target sites in *Tead* genes in vertebrates is required to answer this question.

### A roadmap to phenotypic target identification

Here we provide a unique example of *de novo* miRNA phenotypic target identification in a vertebrate *in vivo* model. To date, a very limited number of studies have identified miRNA phenotypic targets through the deletion of the miRNA binding site in the 3’ UTR region of target mRNAs (9–12), which corresponds to the gold standard of miRNA phenotypic target identification and validation (8). These prior success stories were made possible due to the use of non-vertebrate metazoan models (9–11), existing experimental evidence pointing out the identity of the phenotypic target (9, 11, 12), or the use of an *in vivo* screen in *C. elegans* (10). To our knowledge, a phenotypic miRNA target was never *de novo* (*i.e*., without prior knowledge of its possible identity) identified in a vertebrate *in vivo* model. The identification of biologically relevant miRNA targets is therefore a critical issue, especially in vertebrates, given the high number of possible targets that can be computationally predicted. In this context, our study provides an example of successful miRNA phenotypic target identification. Throughout the process, choices were made that can probably be modulated depending on the biological system used and the outputs of computational analyses. Ultimately, the overall goal of our stepwise approach was to drastically narrow down the list of computationally predicted targets to a manageable short list of possible targets. In our case, we used target identity as well as binding affinity and evolutionary conservation of the miRNA/mRNA interaction to further refine our analysis and select *tead3b* as a credible phenotypic target. Our approach relies on the comparison of miRNA-guided target repression with inter-individual fluctuations in target expression among wild-type specimens. It therefore captures other criteria affecting the amplitude of miRNA-guided repression (number or position of miRNA binding sites in the 3′ UTR, quality of 3′ UTR sequence annotation, or the influence of context sequences): the principle of our method is to favor genes whose biological functionality is most likely to be affected by miRNA-guided repression, rather than to merely select the most efficiently repressed genes. Because our approach relies on gene expression, binding affinity, and sequence conservation features, it is likely to be relevant to any biological model. However, the importance of several key parameters should be emphasized to increase the chances of success of the overall computational screen, as discussed below.

Firstly, it should be reminded that miRNA-regulations make sense only in a cellular context. The use of homogenous biological material is therefore a prerequisite for successful phenotypic identification. While isolated homogenous cell populations would probably be ideal as RNA-seq material, we show here that the isolation of homogenous biological entities (*i.e*., ovarian follicles of a specific size in our case) can be sufficient. Indeed, our data clearly demonstrate that the type (*i.e*., size/stage) of ovarian follicles used has a major impact on the output of the analysis. In a previous study, we used whole ovaries composed of ovarian follicles of many different sizes and searched for possible targets among differentially expressed genes between WT and mutant ovaries after microarray analysis(23). The two possible targets that we discussed, *clockb* (ENSORLG00000004495) and *stat3* (ENSORLG00000004061) are not differentially expressed in the present study in individualized ovarian follicles. Together, these observations clearly illustrate the impact of the type and homogeneity of the biological material used for analysis.

Secondly, our results clearly highlight the importance of using a high number of biological replicates in the RNA-seq data. Because miRNA-triggered gene repression is modest and because gene expression is naturally variable, it is critical to be able to precisely quantify the variability in gene expression in WT and KO individuals using a sufficient number of biological replicates. In the present study, we used 9-11 biological replicates per experimental group. While this could be a limitation in some experimental models, variability in gene expression in WT samples is a critical information that can be inferred, even though less precisely, using a lower number of replicates. Further analyses are needed to estimate the optimum and minimum number of biological replicates required to identify genes that are credible targets based on the variability in their natural expression.

Finally, it clearly appears that a comprehensive understanding of the phenotype is necessary, or at least useful. In our study, we used a quantitative non-ambiguous phenotype (*i.e.*, the number of eggs) that is under miR-202 control. Being able to link this quantitative phenotype to changes in the expression of candidate target genes proved to be extremely useful and most likely significantly contributed to the overall success of our approach.

### The single target paradigm

Here we show that miR-202 is regulating a major organism-level phenotype (*i.e.*, egg production) through a single target within the Hippo pathway. In addition, our data suggest that miR-202 does not regulate egg production through *vgll4b* and *sav1*, two other members of the Hippo pathway. While we cannot rule out the possibility that other targets are also regulating egg production, this result is fully consistent with examples of phenotypic targets that have been reported in Metazoans (9–12) in which a single target is responsible for controlling the phenotype, or at least most of the phenotype. These results therefore contradict the notion that the numerous genes exhibiting miRNA binding sites in their 3′ UTRs coordinately contribute to biological phenotypes (5, 8). Together, these data are consistent with the hypothesis that some miRNAs act, at least when they are regulating a specific phenotype, through a single mRNA target, or a limited number of targets, rather than through multiple targets. They further support the hypothesis that only a limited number of key biological targets would be functionally sensitive to miRNAs (2, 46).

In our previous study (23), we showed that in addition to the reduced egg number phenotype studied here, *mir-202 KO* females also produced eggs exhibiting a lower developmental success. This reduced egg developmental competence resembled a reduced fertilization phenotype even though impaired fertilization was not demonstrated. Interestingly, the *tead3* mutant line described here did not exhibit a decrease in developmental competence (Figure 6C) in contrast to the *mir-202* KO line (Figure 1D). These observations suggest that while miR-202 regulates egg production through *tead3b*, egg developmental competence is regulated through another target, or several other targets, that remain to be identified. In fish, fertilization success relies on the formation of a channel (*i.e*., the micropyle) able to guide the spermatozoa to the oocyte membrane to subsequently allow fertilization. Recent evidence has shown that the micropyle originates from a single cell (*i.e*., the micropylar precursor cell) that differentiates within the granulosa layers under the control of Taz, the main effector of the Hippo pathway (43). While *tead3b* is clearly not responsible for the decrease in fertilization or developmental success observed in *mir-202* KO females, the limited increase in developmental success observed in *tead3b* [-40]UTR mutant females could be the indirect consequence of a perturbation in the Hippo pathway in this model. These data also suggest that miR-202 could be acting on the Hippo pathway through different targets to regulate egg production and egg fertilization. Identifying miR-202 target(s) in a single cell (*i.e*., the micropylar precursor cell) among all ovarian follicular cells is methodologically challenging and will require further investigations.

## Data availability

All transcriptomics datasets generated as part of the study are publicly available. RNA-seq datasets are available on the NCBI SRA portal (https://www.ncbi.nlm.nih.gov/sra) under accession number PRJNA825376.

## Supplementary data

**Table S1.** Normalized RNAseq data at the early-vitellogenic (EV) stage and subsequent successive filters applied.

**Table S2.** Normalized RNAseq data at the late-vitellogenic (LV) stage and subsequent successive filters applied.

**Figure S1.** Conservation of miR-202/*tead3b* pairing geometry among Neopterygii (target-centric view).

## Acknowledgments

The authors thank Guillaume Gourmelen and the INRAE LPGP ISC staff for animal rearing.

