## Supplementary material for "Looking for a needle in a haystack: de novo phenotypic target identification reveals Hippo pathway-mediated miR-202 regulation of egg production": Figure S1

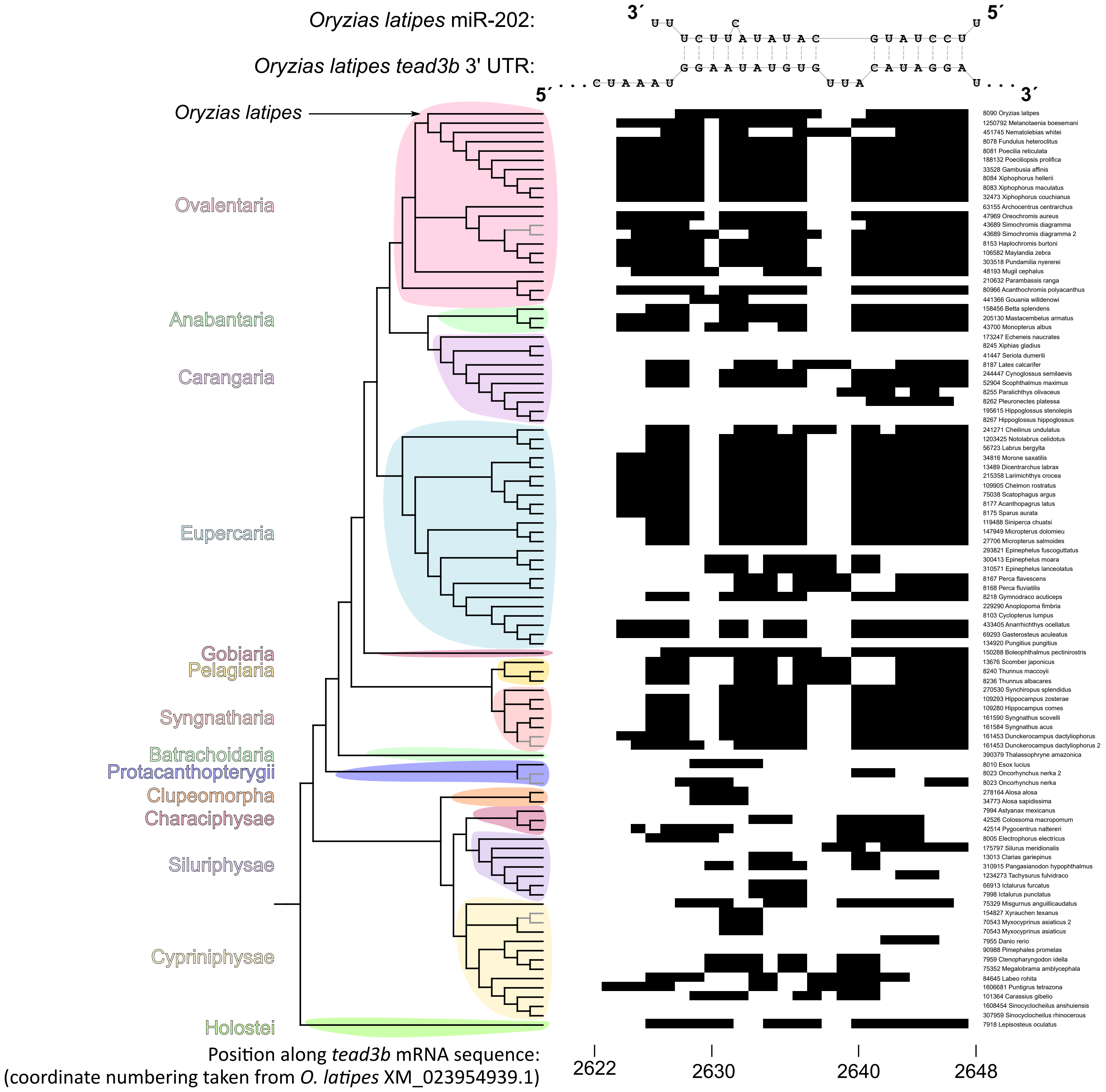


Supplementary figure 1. Conservation of miR-202/*tead3b* pairing geometry among Neopterygii (target-centric view; see Figure 6 for miRNA-centric view of the duplex). Each column in the heatmap represents a *tead3b* mRNA nucleotide (from nt 2622 to 2648 in the *O. latipes* mRNA, and orthologous positions in other species), each row represents one species (phylogenetic cladogram shown on the left margin). Four species possess genes expressing two distinct mature miR-202 sequences: for these species, two alternative pairing geometries are therefore possible between miR-202 and the site in *tead3b* 3´ UTR; they are represented by two rows on the heatmap, and linked by a grey fork in the cladogram. In the heatmap, a black rectangle denotes a paired nucleotide (either a standard Watson-Crick pair, or a wobble G-U pair), and a white rectangle denotes an unpaired nucleotide. *O. latipes* miR-202 and target site sequences are shown at the top for reference.
